# Vascular plants are strong predictors of multi-taxon species richness

**DOI:** 10.1101/252999

**Authors:** Ane Kirstine Brunbjerg, Hans Henrik Bruun, Lars Dalby, Camilla Fløjgaard, Tobias G. Frøslev, Toke Thomas Høye, Irina Goldberg, Thomas Læssøe, Morten D. D. Hansen, Lars Brøndum, Lars Skipper, Kåre Fog, Rasmus Ejrnæs

## Abstract

Plants regulate soils and microclimate, provide substrate for heterotrophic taxa, are easy to observe and identify and have a stable taxonomy, which strongly justifies the use of plants as bioindicators in monitoring and conservation. However, insects and fungi make up the vast majority of species. Surprisingly, it remains untested whether plants are strong predictors of total multi-taxon species richness. To answer this question, we collected an extensive data set on species richness of vascular plants, bryophytes, macrofungi, lichens, plant-galling arthropods, gastropods, spiders, carabid beetles, hoverflies and OTU richness from environmental DNA metabarcoding. Plant species richness per se was a moderate predictor of richness of other taxa. Taking an ecospace approach to modelling, the addition of plant-derived bioindicators revealed 1) a consistently positive effect of plant richness on other taxa, 2) prediction of 12-55% of variation in other taxa and 48 % of variation in the total species richness.

## INTRODUCTION

The majority of species worldwide are still undescribed and nowhere on Earth are all the locally resident species known. Even for the vast majority of known species, their distribution range and population sizes remain unknown. As a result, understanding the causes of spatial variation in biological diversity represents a perpetual challenge for ecological science (Pennisi 2005), with few generally accepted causal mechanisms and models (e.g., Grace *et al*. 2014; Pärtel *et al*. 2016; DeMalach *et al*. 2017). Moreover, we also lack cost-effective, validated methods for assessing biodiversity. Given the global biodiversity crisis, the need for establishing causes for spatial and temporal variation in biodiversity is acute (Hill *et al*. 2016; Ceballos *et al*. 2017).

Brunbjerg *et al*. (2017b) proposed *ecospace* as a unifying framework for assessing and managing variation in biodiversity within regional species pools. *Ecospace* represents the variation in local environment separated into abiotic position (in environmental hyperspace), biotic expansion (diversification of organic matter) and spatio-temporal continuity. Vascular plants are the dominant primary producers of terrestrial ecosystems and plants are quite accurate indicators of the abiotic environment, in which they grow. Here, we test whether plant community composition may be used to predict the overall biodiversity through bioindication of abiotic position and biotic expansion in *ecospace*.

The intractability of total species surveys, has motivated the use of surrogate species in conservation planning (Margules & Pressey 2000; Sarkar & Margules 2002), with the underlying assumption that species richness correlate among taxonomic groups (Gaston 1996). Surrogate species are assumed to reflect the distribution of other species or taxonomic groups, but also to indicate the occurrence of habitats and species of high conservation value (Pearman & Weber 2007). Much research has focused on testing surrogacy and selecting the best taxa (reviewed in Rodrigues & Brooks 2007). It has generally been found that correlations in species richness across taxa vary depending on spatial scale (grain and extent), geographic location (Hess *et al*. 2006) and taxonomic focus (e.g., Wolters *et al*. 2006). Overall, biodiversity surrogacy studies have shown only weak predictive power (Su *et al*. 2004; Rodrigues & Brooks 2007). Similarity in community composition shows more convincing results than species richness. This may be because species composition exhibits a stronger relationship to environmental gradients than does species richness (Su *et al*. 2004; Prober *et al*. 2015). In general, using multi-taxon surrogacy to select areas of conservation interest has been proposed as a more robust measure of biodiversity than single-taxon surrogacy (Smith-Patten & Patten 2015). Environmental characteristics of an area or biotope, i.e. environmental surrogates, have also been tested, but found to be less useful for prediction than cross-taxon surrogates (Rodrigues & Brooks 2007).

Plants are very often included in biodiversity monitoring programs for several good reasons: Plants are sessile and reflect conditions at the place of observation, plants are less seasonal and their detection is less dependent on weather conditions than are fungi and arthropods, plants occur in most ecosystems, and skilled field botanists are generally available. Despite their wide use, the evidence for using plants as surrogates for multi-taxon biodiversity is equivocal (Sætersdal *et al*. 2004; Wolters *et al*. 2006; Myšák & Horsák 2014). Complex metrics representing habitat quality based on weighted measures of vegetation structure (e.g. native plant species richness, number of trees with hollows and total length of fallen logs), plant species richness and functional diversity, have also been suggested to work as surrogates for overall biodiversity, but with limited success (Kwok *et al*. 2011; Hanford *et al*. 2017). Despite the moderate support, plant-based monitoring programs and conservation guidelines remain a common practice, even at supranational levels. For example, in the EU Habitats Directive (1992), plants are implicitly assumed to work as indicators for both habitat types (so-called Annex 1 habitats) and their conservation status. Moreover, averaging plant indicator values (e.g., Ellenberg Indicator Values, Ellenberg *et al*. 1991) is commonly used in vegetation studies to assess local conditions (e.g., Diekmann 2003). The validity of plant-based bioindication has been confirmed by direct measurement of the environmental conditions and by plant growth experiments (e.g., Schaffers & Sýkora 2000; Bartelheimer & Poschlod 2016).

Our approach to bioindication follows the ecospace framework (Brunbjerg *et al*. 2017b). Since plants can be used as indicators of the abiotic environment, they can describe the ecospace *position*. With regard to ecospace *expansion*, i.e. the differentiation of organic matter, each different plant species constitute a potential substrate for specialized insects and fungi (Strong *et al*. 1984; Basset *et al*. 2012; Zhang *et al*. 2016; Brunbjerg *et al*. 2017b). While the species richness responses to ecospace position along environmental gradients may vary among taxonomic groups, we generally expect ecospace expansion by plant species richness to have a positive effect, at least on the richness of heterotrophic taxa.

Plants are highly responsive to land-use change, which usually involves replacement of natural vegetation by crops and weeds, a process generally considered a major cause of biodiversity loss (Pimm *et al*. 2014; Lehsten *et al*. 2015). Effects may be detectable in plant communities for decades or even longer (Gustavsson *et al*. 2007; Hermy & Verheyen 2007). We expect a plant-derived land-use intensity indicator to be useful for prediction of multitaxon species richness, especially when used in combination with a plant-derived indicator of abiotic conditions and plant species richness.

Here we put the value of a plant species list to a test. We use a comprehensive dataset of 130 sites, each 40 × 40 m, sampled for richness of plants and a range of other taxa including DNA derived OTUs (Operational Taxonomic Units), collectively spanning the major environmental variation in terrestrial habitats within a region (from wet to dry, nutrient rich to nutrient poor, early to late succession). These data allow us to investigate the following questions:

1. Can plant species richness be used as surrogate for species richness of other taxa across habitat types?
2. Does plant-inferred *ecospace*, in the form of a combination of plant species richness and environmental bioindication, improve the prediction of species richness of other taxa?

Our study is the first to comprehensively validate the ubiquitous use of plants in conservation planning and monitoring, which has been incorporated – based on anecdotal evidence and tradition – into national and supranational legislation.

## METHODS

We selected 130 study sites (40 m × 40 m) evenly distributed across five geographic regions in Denmark (Fig. S1 in Supporting Information). Within each region, sites were placed in three clusters for logistical reasons, but with a minimum distance of 500 m between sites to reduce spatial covariance. Site selection was stratified according to primary environmental gradients. We allocated 30 sites to cultivated habitats and 100 sites to natural habitats. The cultivated subset was stratified according to major land use type and the natural subset was selected amongst uncultivated habitats and stratified according to gradients in soil fertility, soil moisture and successional stage. We deliberately excluded saline and aquatic habitats, but included temporarily inundated depressions as well as mires and fens. The final set of 24 habitat strata consisted of the following six cultivated habitat types: Three types of fields (rotational, grass leys, set aside) and three types of plantations (beech, oak, spruce). The remaining 18 strata were natural habitats, constituting all factorial combinations of: Fertile and infertile; dry, moist and wet; open, tall herb/scrub and forest. These 24 strata were replicated in each of the five geographical regions. We further included a subset of 10 perceived hotspots for biodiversity in Denmark, selected subjectively by public voting among active naturalists in the Danish conservation and management societies, but restricted so that each region held two hotspots. See Brunbjerg *et al*. (2017a) for more details on site selection and stratification.

### Collection of biodiversity data

The field inventory aimed at an unbiased and representative assessment of the multi-taxon species richness in each of the 130 sites. We collected data on vascular plants, bryophytes, lichens, macrofungi, arthropods and gastropods. Due to the limited size of each site, we excluded vertebrates from consideration. We conducted a complete inventory of vascular plants and bryophytes. For the remaining taxa, which are more demanding to find, catch, and identify, we aimed at collecting a reproducible and un-biased sample through a standardized effort. Each site was carefully examined for gastropods, lichens, plant-galling arthropods (a single visit per group) and macrofungi (three visits at different times within the autumn season), actively searching contrasted microhabitats and substrates (soil, herbaceous vegetation and debris, wood, stone surfaces and bark of trees up to 2 m). Similarly, a standard set of passive traps was used to survey insects (pitfall traps, meat-baited and dung-baited traps, yellow pan traps and Malaise traps) during periods of standard length and timing. Biodiversity survey methods are detailed in (Brunbjerg *et al*. 2017a).

### OTU richness from DNA metabarcoding

Massive parallel sequencing of amplified marker genes from environmental DNA from e.g., a soil sample – also known as DNA metabarcoding – is increasingly used to assess biological communities (Taberlet *et al*. 2012). Sequences are traditionally grouped into ecologically meaningful Operational Taxonomic Units (OTUs), which are then used as richness estimates (Bálint **et al*.* 2016; Frøslev *et al*. 2017). In order to extend our biodiversity assessment to organisms assumed to be poorly represented in traditional surveys, we included OTU richness of fungi and general eukaryotes from soil samples and of arthropods from Malaise traps (called fungal OTUs, eukaryote OTUs and Malaise OTUs, respectively). Taxa preferentially caught in Malaise traps (mostly Diptera and Hymenoptera) remain largely unidentified, but Malaise OTUs may have a minor overlap with respect to the hoverfly and spider specimens caught in these traps as these were pooled with contents of the other traps in richness estimates for these two groups.

We collected soil from all sites and subjected it to metabarcoding through DNA extraction, PCR amplification of genetic marker regions (DNA barcoding regions) and massive parallel sequencing on the Illumina platform as described in (Brunbjerg **et al*.* 2017a). The soil sampling scheme included the mixing of 81 soil cores from each site in an attempt to get a representative sample. For this study, we used sequencing data from genes amplified with primers targeting fungi and eukaryotes. For eukaryotes, we used the primers 18S_allshorts (Guardiola *et al*. 2015, 2016) with a slight modification of the forward primer (TTTGTCTGGTTAATTCCG) to exclude fungi. For fungi, we amplified the ITS2 region with primers gITS7 (Ihrmark *et al*. 2012) and ITS4 (White *et al*. 1990). Furthermore, we extracted DNA from the ethanol of Malaise traps and subjected it to sequencing. For this we amplified a region of the CO1 gene of primarily arthropods with primers ZBJ-ArtF1c and ZBJ-ArtR2c (Zeale *et al*. 2011), and subjected it to a sequencing approach similar to the other markers. We extracted, amplified, and sequenced DNA from two independent trapping periods for each site and combined these to obtain a single richness measure.

The bioinformatic processing of the sequence data followed the strategy outlined in (Brunbjerg **et al*.* 2017a) including post-clustering curation with the LULU algorithm (Frøslev **et al*.* 2017) in order to obtain reliable alpha diversity estimates for OTU data. Although, it is widely acknowledged (e.g., Bálint **et al*.* 2016) that species richness is difficult to estimate from sequencing data of environmental DNA, Frøslev **et al*.* (2017) showed that careful bioinformatics processing can produce richness measures based on OTU data with strong correlation to richness metrics based on survey data. For this study, a simple OTU count was used as a DNA based richness metric, after ensuring that variation in sequencing depth between samples only had a minor impact (results not shown).

### Conservation Index

A metric of conservation value was produced to test if plants can predict the richness of species of conservation concern. For vascular plants, macrofungi, lichens, gastropods, spiders and arthropods we used the national red list for Denmark (Wind & Pihl 2004). For taxonomic groups lacking a national red list (bryophytes and galling arthropods) an expert-based red listing was created for this project using the same criteria as the official red lists (bryophyte expert: Irina Goldberg, galling arthropod expert: Hans Henrik Bruun). Each red listed species contributed to a weighted score of threatened species per site (the Conservation Index) as follows: red list status RE (regionally extinct) and CR (critically endangered) = 4 points, red list status EN (moderately endangered) = 3 points, red list status VU (vulnerable) = 2 points, and red list status NT (near threatened) and DD (data deficient) = 1 point.

### Abiotic Factors and Environmental Calibration

We used field-measured abiotic variables to validate the plant-based environmental calibration. Environmental recordings and estimates included soil pH, soil C, N and P, soil moisture, leaf N and P concentrations, air temperature, light intensity, soil surface temperature, and humidity, number of trees >40dbh, dead wood volume, and vapor pressure deficit (VPD). For further details on methods for collection of the abiotic data see (Brunbjerg **et al*.* 2017a).

Mean Ellenberg Indicator Values were calculated for light conditions, soil nutrient status, soil pH and soil moisture based on the plant lists for each site and the species’ abiotic optima (Ellenberg *et al*. 1991).

### Natural Habitat Index

To supplement the plant-based environmental calibration, we calculated a plant-based natural habitat index reflecting land use intensity using 115,071 vegetation quadrats from the national monitoring of terrestrial biodiversity (Svendsen *et al*. 2005; Nielsen *et al*. 2012). Vegetation data were grouped in quadrats from Annex 1 habitats (A1) (EU Habitats Directive 1992) of conservation value (excluding nitrophilous tall herb fringes), other quadrats sampled in natural areas (Na), and quadrats sampled in agricultural (Ag) landscapes (road verges, hedges, soil banks etc.). For each species in the dataset a natural habitat score was calculated as:

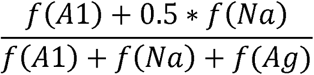

Where f () = frequency of species in the mentioned habitat category.

The species level score is thus a number between 0 and 1, where 0 implies that the species only occurs in agricultural biotopes and 1 implies that the species only occurs in habitats of conservation concern. The natural habitat index was calculated for each site as the mean of species scores. It reflects land use and land use history under the assumption that protected natural areas have been less intensively managed than farmland habitats.

### Analyses

We used Spearman rank correlation to test for correlations between species richness of vascular plants and the richness of other taxonomic groups including macrofungi, lichens, bryophytes, gastropods, plant galling arthropods, carabid beetles, hoverflies, spiders, fungal OTUs, eukaryote OTUs and Malaise OTUs. To validate the plant-based environmental calibration, Spearman correlations were calculated between Ellenberg Indicator Values, the natural habitat index, and measured environmental variables (soil moisture, soil C, N, and P, leaf N, and P, soil pH, surface, and air temperature, light intensity, number of trees >40dbh, dead wood volume, and vapor pressure deficit (VPD)).

We also grouped the 130 study sites into five different land use intensity categories from protected Annex 1 habitats, over other uncultivated areas, plantation forest and extensively farmed habitats to intensive farmland. ANOVA followed by Tukey’s post hoc tests was used to test for differences in mean natural habitat index value between the five habitat types.

To assess the efficiency of plants as indicators for other taxonomic groups, we performed multiple regression with species richness of macrofungi, bryophytes, lichens, plant galling arthropods, gastropods, carabid beetles, hoverflies, spiders, the Conservation Index, soil fungal and eukaryote OTU richness and Malaise OTU richness as response variables and plant species richness and plant-derived bioindicators as explanatory variables. Data exploration was carried out following the protocol described in Zuur *et al*. (2010). Collinearity was assessed using Variance Inflation Factors (VIF) sequentially disregarding variables showing VIF values >3 from the VIF calculations (Zuur *et al*. 2010). Ellenberg nutrient status was found to be correlated with Ellenberg pH and our Conservation Index [VIF >3] and was excluded from all models. If GAM smoothers fitted to the residuals of the models were conservatively significant (p < 0.01) for any of the predictors, we included polynomials in the final model. To account for the geographically nested design, we used Generalized Linear Mixed Modelling (GLMM) with a Poisson distribution and a log link function and with cluster (n=15) as random intercept. If overdispersion was detected, we used GLMM with Negative Binomial error distribution instead of Poisson error distribution (Hilbe 2011), given that the Deviance Information Criterion (DIC) indicated a better fit based on a ΔDIC < 2, criterion. Model assumptions were verified by plotting residuals versus fitted values and versus each covariate in the model. We assessed the residuals for spatial dependency. Modelling was performed in R version 3.4.3 (R Core Team 2017), models were fitted by approximate Bayesian inference using the INLA package (Rue *et al*. 2009). Explanatory variables were scaled prior to model implementation. Marginal posterior distributions were summarized by 95 % Bayesian credible intervals corresponding to the 0.025 and 0.975 quantiles of the posterior distribution (Zuur *et al*. 2017). We chose not to perform model selection, but covariates can be considered statistically important if the 95% Bayesian credible intervals (BCI) do not overlap zero (Zuur *et al*. 2017). As an aid in the interpretation and comparison of model-based predictions, we calculated Pearson’s product moment correlations between fitted values (only for fixed variables) and observed species richness of the response variables (pseudo R^2^).

## RESULTS

### Biodiversity data

The total number of species of plants, macrofungi, bryophytes, lichens, plant galling arthropods, gastropods, carabid beetles, hoverflies and spiders per site ranged from 78 to 481. Species number per site ranged from 11-134 for plants (with up to 5 red-listed plant species per site), 0-24 for gallers (up to 3 expert-assessed red-listed gallers per site), 0-180 for macrofungi (up to 15 red listed macrofungi species per site), 0-33 for lichens (up to 12 red-listed lichen species per site), 0-50 for bryophytes (up to 3 expert-assessed red-listed bryophyte species per site), 7-53 for spiders (up to 8 red-listed spider species per site), 0-21 for carabid beetles (up to 4 red-listed carabids per site), and 0-21 for hoverflies (up to 2 red-listed hoverflies per site). After bioinformatic processing the soil fungal OTU dataset resulted in an OTU richness per site ranging from 66 to 476. The soil eukaryote OTU data had a richness per site ranging from 206 to 1549. The Malaise OTU dataset had a richness per site ranging from 25 to 160.

### Plant-derived environmental bioindication

Plant-derived environmental bioindication was supported by correlation to independent data. Community mean Ellenberg Indicator Values correlated well with corresponding measured abiotic factors (Table S1 in Supporting Information). The natural habitat index scored highest for EU Habitats Directive Annex 1 habitats, intermediate for other uncultivated habitats, plantation forests and extensively farmed sites, and lowest for sites with intensive cropping (Fig. S2). The natural habitat index correlated negatively with Ellenberg nutrient status, indicating that plants with affinity to natural habitats generally occur in infertile environments (Fig. S3). Plant species richness correlated positively with plant-derived Ellenberg soil pH and soil nutrient status and with measured soil pH, C, N, and P. In contrast, there was a negative correlation between plant species richness and the number of trees > 40dbh, dead wood volume, and canopy height (Table S1).

### Plant species richness as surrogate for other taxonomic groups

Spearman rank correlation between plant richness and species richness for other taxa revealed no significant correlation for macrofungi, bryophytes, lichens, carabid beetles and summed richness. The richness of plant-galling arthropods, gastropods, spiders and hoverflies showed significant positive relationships with plant richness (Table S2).

### Plant-derived bioindication of species richness

Plant species richness was important for the prediction of species richness of all surveyed taxonomic groups except carabid beetles (Table 1, Fig. 1). Plant species richness was also important for richness of fungal OTUs, Malaise OTUs and for the Conservation Index (Table 1, Fig. 1). In all cases, except carabid beetles, the relationship between plant species richness and other groups was positive (Fig. 1).

**Figure 1.**
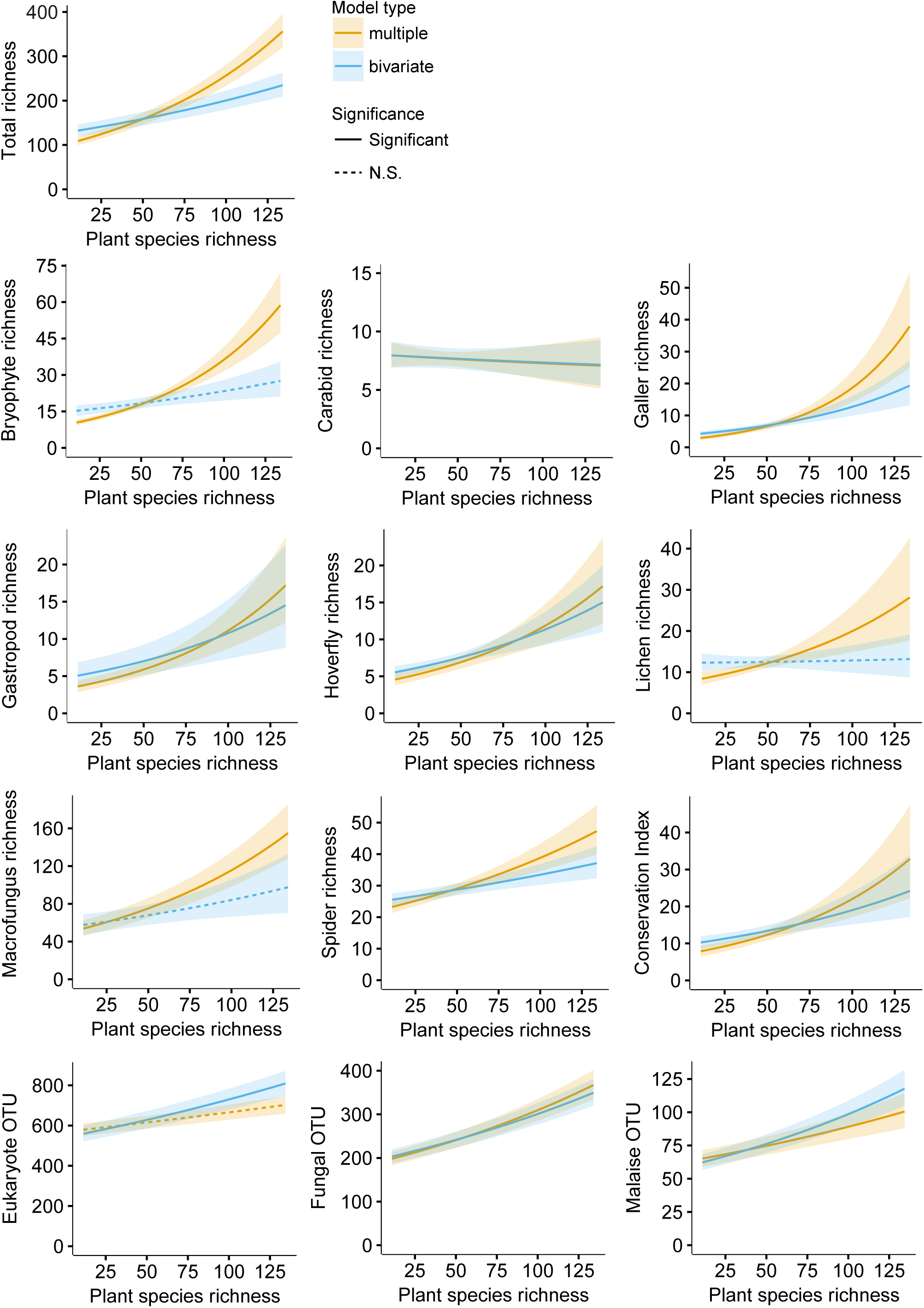
Relationships between plant species richness and species richness of other taxonomic groups, Conservation Index, and soil fungal, eukaryote and Malaise OTU richness in multiple and bivariate regressions and their 95% BCI. Stippled versus full line indicates significance and non-significance at the 0.05 level (parameter estimates whose 95% BCI did not overlap zero), respectively. For model details see Table 1.

**Table 1.** Model results for the full GLMM models (Poisson or Negative Binomial). Model results show the effects of plant species richness (plant_rich), Ellenberg light (E_light), Ellenberg pH (E_pH), Ellenberg moisture (E_moisture) and natural habitat index (nat_index), on species richness of various taxonomic groups (total, bryophytes, carabids, gallers, gastropods, hoverflies, lichens, macrofungi, and spiders), Conservation Index, and OTU richness (eukaryote, fungal, and Malaise). Intercept, parameter estimates (marginal posterior means), and 95% Bayesian credible intervals (BCI, i.e. the 0.025 and 0.975 quantiles of the posterior distribution) are given in parentheses. Parameter estimates with 95% BCI not overlapping zero are shown in bold.

| Response | Intercept | plant_rich | E_light | E_moisture | E_pH | nat_index | nat_index<sup>2</sup> |
| --- | --- | --- | --- | --- | --- | --- | --- |
| Total richness | **5.00**<br>(4.95, 5.06) | **0.22**<br>(0.15, 0.29) | **-0.33**<br>(-0.39, -0.27) | **0.11**<br>(0.06, 0.17) | **-0.13**<br>(-0.22, -0.05) | 0.04<br>(-0.04, 0.12) | - |
| Bryophytes | **2.78**<br>(2.7, 2.86) | **0.35**<br>(0.24, 0.45) | **-0.38**<br>(-0.48, -0.28) | **0.24**<br>(0.16, 0.32) | **-0.36**<br>(-0.5, -0.23) | **0.14**<br>(0.02, 0.27) | - |
| Carabids | **2.05**<br>(1.96, 2.13) | -0.03<br>(-0.14, 0.08) | 0.02<br>(-0.07, 0.11) | 0.02<br>(-0.07, 0.11) | 0.06<br>(-0.08, 0.20) | **-0.12**<br>(-0.24, -0.01) | - |
| Gallers | **1.74**<br>(1.61, 1.87) | **0.55**<br>(0.38, .73) | **-0.53**<br>(-0.69, -0.38) | 0.08<br>(-0.05, 0.21) | -0.20<br>(-0.41, 0.01) | 0.01<br>(-0.19, 0.22) | - |
| Gastropods | **1.70**<br>(1.58, 1.81) | **0.29**<br>(0.15, 0.43) | **-0.72**<br>(-0.86, -0.6) | **0.29**<br>(0.17, 0.41) | **0.29**<br>(0.12, 0.47) | -0.02<br>(-0.19, 0.16) | - |
| Hoverflies | **1.87**<br>(1.75, 1.98) | **0.27**<br>(0.13, 0.42) | **0.38**<br>( 0.25, 0.52) | **0.24**<br>(0.13, 0.35) | **-0.20**<br>(-0.38, -0.02) | **-0.26**<br>(-0.43, -0.10) | - |
| Lichens | **2.42**<br>(2.28, 2.57) | **0.26**<br>(0.07, 0.45) | **-0.49**<br>(-0.69, -0.31) | **0.15**<br>(0.01, 0.29) | -0.23<br>(-0.46, 0.01) | **0.42**<br>(0.17, 0.68) | - |
| Macrofungi | **4.30**<br>(4.17, 4.42) | **0.17**<br>(0.05, 0.29) | **-0.43**<br>(-0.56, -0.31) | 0.04<br>(-0.06, 0.13) | -0.10<br>(-0.25, 0.04) | 0.11<br>(-0.05, 0.27) | **-0.26**<br>(-0.34, -0.17) |
| Spiders | **3.33**<br>(3.28, 3.38) | **0.14**<br>(0.08, 0.2) | 0.01<br>(-0.04, 0.07) | **0.09**<br>(0.04, 0.13) | **-0.13**<br>(-0.21, -0.06) | **-0.07**<br>(-0.14, 0.00) | - |
| Cons. Index | **2.43**<br>(2.3, 2.56) | **0.26**<br>(0.11, 0.42) | **-0.46**<br>(-0.62, -0.3) | 0.03<br>(-0.1, 0.16) | 0.02<br>(-0.18, 0.22) | **0.66**<br>(0.44, 0.89) | - |
| Eukaryote OTU | **6.41**<br>(6.36, 6.47) | 0.04<br>(-0.04, 0.11) | 0.01<br>(-0.05, 0.08) | **0.13**<br>(0.07, 0.19) | **0.10**<br>(0.00, 0.19) | -0.06<br>(-0.14, 0.02) | - |
| Fungal OTU | **5.45**<br>(5.40, 5.51) | **0.15**<br>(0.07, 0.22) | **-0.09**<br>(-0.16, -0.03) | **-0.06**<br>(-0.12, 0.00) | -0.01<br>(-0.10, 0.08) | -0.05<br>(-0.13, 0.03) | - |
| Malaise OTU | **4.30**<br>(4.24, 4.37) | **0.10**<br>(0.03, 0.18) | **0.13**<br>( 0.06, 0.19) | 0.00<br>(-0.06, 0.06) | 0.05<br>(-0.05, 0.14) | -0.03<br>(-0.12, 0.05) | - |

Multiple regression of species richness of the selected taxa varied in percent explained variation by fixed variables from 12 % for carabid beetle richness to 55 % for gastropod species richness (Fig. 2). The corresponding bivariate regression between plant richness and other richness metrics explained below 5 % of variation in total richness for gastropods, total richness, bryophytes, fungi and eukaryote OTU richness, 5-10% explained variation for hoverflies, spiders and Conservation Index and 10-16% for fungal OTU richness, malaise OUT richness and galling insects (Fig 2).

**Figure 2.**
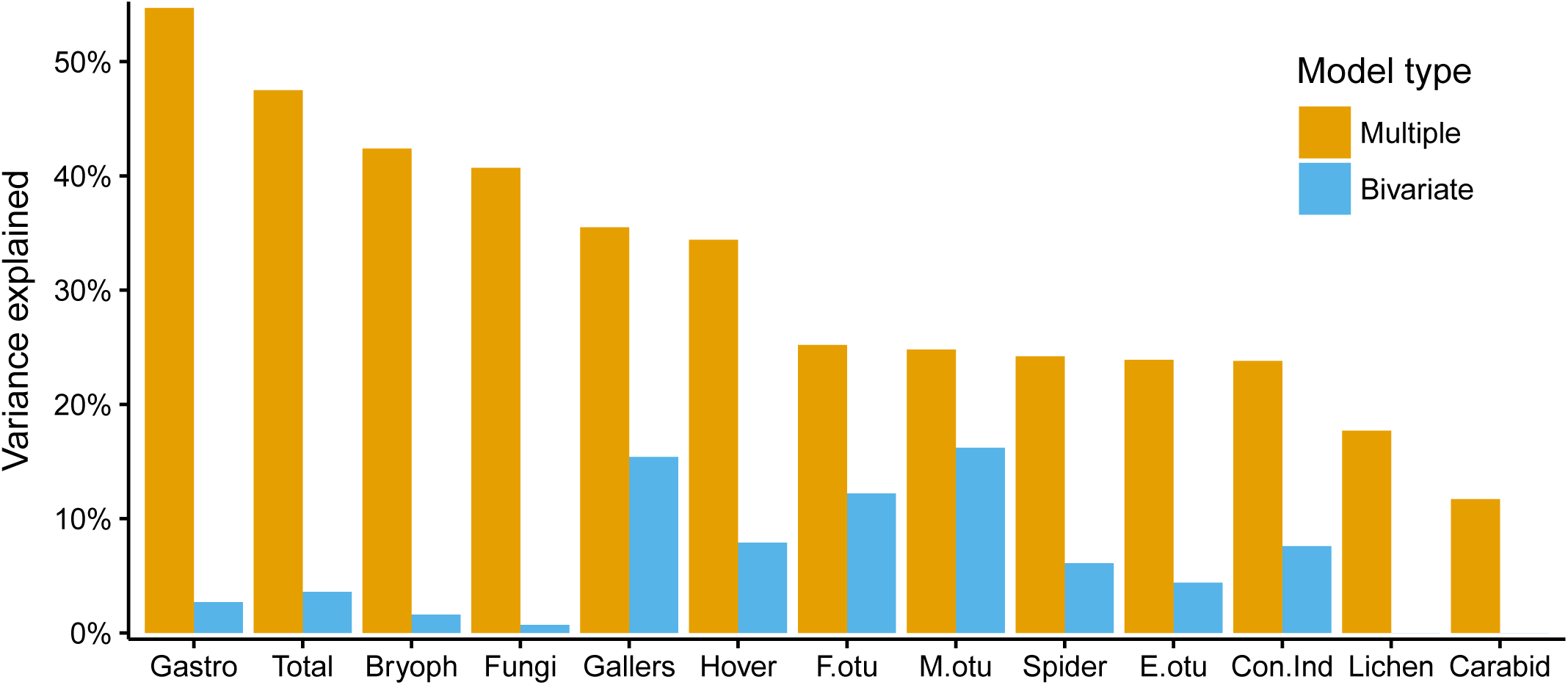
Barplot of percentage variance explained by the multiple and bivariate regressions of species richness of the taxonomic groups, Conservation Index, and soil fungal, eukaryote and Malaise OTU richness. For model details see Table 1. Gastro = Gastropods, Bryoph = Bryophytes, Hover = Hoverflies, F.otu = soil fungal OTU richness, E.otu = soil eukaryote OTU richness, M.otu = Malaise OTU richness, Con.Ind. = Conservation Index.

Ellenberg light was generally important with positive effects for flying insects such as hoverflies and Malaise OTU richness and otherwise negative or neutral effects on species richness and Conservation Index. Increasing Ellenberg moisture seemed to promote the richness of spiders, bryophytes, macrofungi, gastropods, lichens, hoverflies and eukaryote OTU richness, whereas the effect on fungal OTU richness was weak and negative. Ellenberg pH had negative effects on spiders, bryophytes, hoverflies, and total richness, and positive effects on gastropods and eukaryote OTU richness. The natural habitat index had a positive effect on bryophyte and lichen species richness and the Conservation Index, a negative effect on hoverflies, carabids and spiders and a unimodal effect on macrofungi species richness (Table 1, Fig. S4).

## DISCUSSION

Terrestrial biodiversity of heterotrophic organisms relies on the build-up and diversification of organic matter produced by plants. Therefore, it may seem reasonable to use the species richness of vascular plants as a proxy of total biodiversity in science and in conservation planning and management. However, neither plants nor other single taxa have hitherto been confirmed as reliable surrogates for other taxa. This point was supported by our simple cross-taxon correlations. Although plant species richness did correlate positively with four out of eight surveyed taxonomic groups, the correlations were generally weak and the overall performance of vascular plants as biodiversity surrogate was poor. A multivariate modelling approach, with simultaneous inclusion of plant-derived bioindication and plant species richness, showed a much stronger - and consistently positive - effect of plant species richness on the species richness of other taxa, as well as on the Conservation Index and OTU richness measures, except soil eukaryote OTU richness and carabid species richness (Fig. 1).

In monitoring programs, plants are often used as general indicators of conservation status of habitats without explicit testing. While plant species richness in itself may be a poor indicator for the richness of other species groups, plant indication may be a cost-effective approach to estimate environmental conditions (e.g., Diekmann 2003) and possibly also the habitat quality (Andersen *et al*. 2013). We used Ellenberg Indicator Values for light conditions, soil moisture, nutrient conditions and soil pH (Ellenberg *et al*. 1991), which are available only for the Central European flora. Our approach may still be applicable in other parts of the world because species scores from ordination of large and representative vegetation datasets typically reflect major environmental gradients (e.g., Ejrnæs *et al*. 2002) and may replace Ellenberg Indicator Values in much the same way. While bioindication of environmental conditions is well developed, there is currently no standard approach to estimation of habitat quality by plant lists, despite the scientific evidence that plants reflect land-use intensity and land-use history (Hermy & Verheyen 2007). Plant based habitat quality scores may be obtained e.g. by expert judgment (Kowarik 1990) or by empirical evidence (Ejrnæs *et al*. 2002; Ejrnæs *et al*. 2008). In this study, we calculated a naturalness index reflecting the affinity of plants with protected habitats and found that the index correlated closely with Ellenberg nutrient status.

Since an estimated > 85 % of the World’s terrestrial species remain undescribed (Mora *et al*. 2011), choosing surrogates that actually work and reflect general biodiversity is highly relevant. An obvious challenge is that different taxonomic or functional groups respond differently to habitat conditions. For example, lichens and bryophytes growing under extremely infertile conditions, often directly on stone and trees, will show a markedly different richness optimum along a fertility gradient than more competitive vascular plants. Likewise, generalist predatory beetles and spiders may be expected to respond differently than specialist herbivores such as gall wasps or aphids. Therefore, finding that plant species richness, after accounting for the abiotic environment, had a positive effect on species richness of all other taxa except carabids, lends strong support for the idea of biodiversity surrogacy and for vascular plants as optimal surrogates.

The taxonomic groups used in this study varied strongly in their ecological dependence on plants. Plant galling arthropods depend directly on specific plant species as hosts and represent a megadiverse group of phytophagous insects and mites with pronounced host specificity (Jaenike 1990). Hoverflies are generally less host dependent as larvae, but utilize plants as sources of pollen and nectar in the adult life stage. On the other hand, we would expect generalist herbivores or predators such as gastropods, carabid beetles and spiders, as well as primary producers such as lichens and bryophytes, to be causally unrelated to specific plant species. Still, such species may respond to environmental conditions also influencing plant species richness. Macrofungi constitute several functional groups including both generalist decomposers and mycorrhizal symbionts, some of which are specialized on a single plant genus or species. Many decomposer fungi are also specific to certain plant genera or species, while some are necrophagous and highly specialized on arthropods or other fungi. Despite the difference in plant species specificity, the positive effect of plant species richness was consistent across taxonomic groups, pointing to a general applicability of plants as surrogates, even for predatory and decomposer organisms. Plant richness and environmental calibration obtained through bioindication together could account for 48% of the variation in richness of all other surveyed taxa combined. The figures for predicted OTU richness were also supportive with 24-30 % of variation explained.

The amount of explained variation was lowest for species richness of carabid beetles (12%), lichens (18%) and spiders (24%), possibly indicating that vegetation structure or microclimatic properties unrelated to plant community composition may be more important to species in these groups. Mobile generalist predators such as spiders and carabid beetles may rely less on site conditions than sessile species such as plants and fungi. A large proportion of lichens are epilithic or epiphytic on boulders and trees, and therefore partly uncoupled from the prevailing environmental site conditions as reflected by vascular plants. Despite the general usefulness of plants as surrogates, the amount of unexplained variation for specific groups such as lichens, carabids and hoverflies demonstrate that surrogates and indicators should be selected with due reference to spatial scale and the ecology of the target species groups (Zurlini & Girardin 2008; Kwok *et al*. 2011).

In order to test the generality of vascular plants as surrogates, we also included three richness metrics derived from DNA metabarcoding – soil fungal and soil eukaryote OTUs from eDNA and aerial arthropod OTUs from Malaise trap DNA. Despite a thorough sample of 81 regularly spaced soil cores, we have merely covered an approximate 0.01 % of the soil surface of the study sites. This could pose a bottleneck for getting a representative sample of OTUs in diverse and heterogeneous habitats. We assume that the eukaryotic and the fungal genetic markers are targeting a soil community depending on micro-climate and soil composition, and less on vegetation – at least compared to the organisms recorded above ground. Furthermore, the inherent problems in getting reliable richness estimates from eDNA sequencing are widely acknowledged (e.g., Bálint *et al*. 2016). We find it encouraging that the general pattern of a positive effect of plant species richness was reproduced for OTU-richness, albeit insignificant for eukaryotes, and that the explained variance by multiple regressions with plant-derived environmental variables approached 25 % for three taxonomically very different OTU taxa.

Rare and threatened species are particularly important to conservation, and we demonstrated that a plant-based model could explain 23% of the variation in our Conservation Index based on occurrence of red-listed species. Our sites were only 40 m × 40 m and, thus, too small for a representative sampling of very rare species. With larger plots we would expect a higher proportion of explained variation. A general index of site uniqueness could replace the use of rare species for assessment of conservation value of such small sites. Our natural habitat index was the strongest predictor of variation in the Conservation Index which is in accordance with evidence for the preferences of threatened species for rare natural habitats (Pearman & Weber 2007 and references therein).

We find it encouraging that the richness of vascular plants is a consistent positive predictor of multiple functional groups comprised by our multi-taxon species richness estimate. However, looking at the direct trophic effects, we see opportunities for further improvement of plant surrogacy. It has long been acknowledged that plants serve as mutualistic partners for other organisms (e.g., Elton 1949). With respect to the diversification of organic matter, Southwood (1961) and later work by Brändle and Brandl (2001) quantified the richness of phytophagous insects on European trees and showed that the size of their associated biotas vary enormously and predictably, i.e., large, long-lived and omnipresent species may harbor a more diverse pool of insects than small annuals or uncommon species. A thorough examination of reported interactions between plants and associated invertebrates and fungi may be used to create a more powerful surrogate for total biodiversity than the mere number of plant species.

Vascular plants play an important role in the conservation prioritization and monitoring. In this study, we demonstrate that plant species are useful surrogates for biodiversity at large, but only when environmental bioindication is taken into account. Our results support the *ecospace* framework for biodiversity, implying that future research into the diversification of organic matter may further improve the value of plant-related indicators as surrogates of biodiversity in general.

## ACKNOWLEDGEMENTS

RE, LB, IG, TL, TGF, CF and AKB were supported by a grant from VILLUM foundation (Biowide, VKR-023343). We thank Aimee Classen and Greg Newman for assistance with the analysis of soil properties, Vagn Alstrup (†), Ulrik Søchting and Roar Skovlund Poulsen for lichen surveying, Karl-Henrik Larsson for aid in identifying critical corticioid fungi, Leif Örstadius for identifying *Psathyrella* collections. We thank Lars Dyhrberg Bruun for identifying spiders, Monica Oyre for identifying hover flies, and Oskar Liset Pryds Hansen, and Emil Skovgaard Brandtoft for identifying carabid beetles. We also thank a large group of volunteers for assistance during species surveys.

## SUPPORTING INFORMATION

Table S1 - Spearman rank correlations between plant species richness (plant_rich) and plant-derived environmental bioindication (Ellenberg Indicator Values) and measured abiotic factors.

Table S2 - Spearman rank correlation between plant species richness (plant_rich) and species richness of other taxonomic groups and OTU richness.

Figure S1 - Map of Denmark showing the location of the 130 sites grouped into 15 clusters within five regions.

Figure S2 - Boxplot of natural habitat index for sites of five different habitat types.

Figure S3 - Correlation between natural habitat index and Ellenberg nutrient status.

Figure S4 - Relationships between explanatory variables and species richness of various taxonomic groups, Conservation Index and OTU richness.

